## Supplemental Figure 1 for "Breaking the Crosstalk of the Cellular Tumorigenic Network in NSCLC by a Highly Effective Drug Combination"

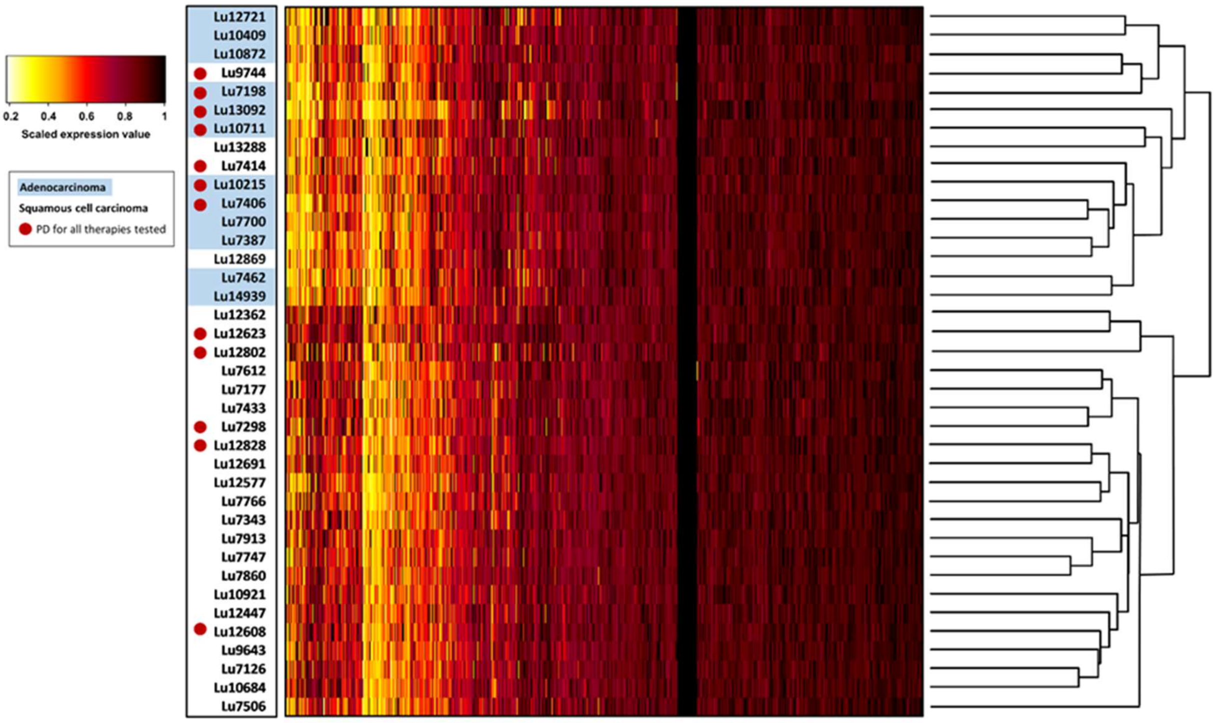


**Supplement Figure 1**

The heatmap represents the relative expression of 19096 HGNC-annotated genes in 38 NSCLC tumors from the EPO tumor biobank. NSCLC tumors with progressive disease (PD) as best response evaluated in patient-derived xenografts are indicated and highlighted with a red circle. Hierarchical clustering of the expression data results in two main clusters reflecting the histology-based gene expression of adenocarcinoma and squamous cell carcinoma PDX.
